# Extracellular ATP plays an important role in systemic wound response activation

**DOI:** 10.1101/2022.01.06.475278

**Authors:** Ronald J Myers, Yosef Fichman, Gary Stacey, Ron Mittler

## Abstract

Mechanical wounding occurs in plants during biotic (*e*.*g*., herbivore or pathogen attack) or abiotic (*e*.*g*., wind damage or freezing) stresses and is associated with the activation of multiple signaling pathways. These initiate many wound responses at the wounded tissues, as well as trigger long-distance signaling pathways that activate wound responses in tissues that were not affected by the initial wounding event (termed ‘systemic wound response’). Among the different systemic signals activated by wounding are electric signals, calcium and reactive oxygen species (ROS) waves, and different plant hormones such as jasmonic acid. The release of glutamate from cells at the wounded tissues was recently proposed to trigger several different systemic signal transduction pathways via glutamate-like receptors (GLRs). However, the role of another important compound released from cells during wounding (*i*.*e*., extracellular ATP; eATP) in triggering systemic responses is not clear. Here we show that eATP that accumulates in wounded leaves and is sensed by the purinoreceptor kinase P2K is required for the activation of the ROS wave during wounding. Application of eATP to unwounded leaves triggered the ROS wave, and the activation of the ROS wave by wounding or eATP application was suppressed in mutants deficient in P2K (*i*.*e*., *p2k1-3, p2k2*, and *p2k1-3p2k2*). In addition, the expression of several systemic wound response transcripts was suppressed in mutants deficient in P2K during wounding. Our findings reveal that in addition to sensing glutamate via GLRs, eATP sensed by P2Ks is playing a key role in the triggering of systemic wound responses in plants.

**One sentence summary:** Extracellular ATP plays an important role in triggering the ROS wave and systemic transcriptomics responses during wounding.

## INTRODUCTION

Mechanical wounding is associated in plant cells with the disruption of cellular integrity and the breakdown of different membrane structures (Farmer et al., 2020; Ikeuchi et al., 2020; Vega-Muñoz et al., 2020). In many cases this disruption results in the rapid and uncontrolled partition of cellular compounds, membrane depolarization, altered pH levels, and/or formation of reactive oxygen species (ROS; Farmer and Goossens, 2019; Shao et al., 2020; Vega-Muñoz et al., 2020). Mechanical wounding can occur in plants during herbivory, wind damage, pathogen infection, or even as part of different abiotic stresses such as severe drought, freezing, heat, or salt stresses that cause the collapse of different subcellular structures. In some cases, wounding is an early response to stress (*e*.*g*., during herbivory), while in others (*e*.*g*., following the activation of programmed cell death in response to pathogen attack) it could be a late event. One of the key processes associated with wounding is the disruption of plasma membrane (PM) integrity and the release of different cellular compounds such as the amino acid glutamate (Glu), or the nucleotide adenosine triphosphate (ATP) that becomes extracellular ATP (eATP; Roux, 2014; Toyota et al., 2018; Shao et al., 2020). Some of the compounds released from cells during wounding are thought to be sensed by different receptors (*e*.*g*., the purinoreceptor kinase P2K that senses eATP; Choi et al., 2014; Chen et al., 2017; Pham et al., 2020), and/or channels (*e*.*g*., the glutamate-like channels GLR3.3 and GLR3.6 that sense Glu; Mousavi et al., 2013; Toyota et al., 2018; Shao et al., 2020; Tian et al., 2020) and trigger wound-induced signal transduction pathways (Monshausen et al., 2009; Tian et al., 2020; Moore et al., 2021). Wound-induced depolarization of the PM is also thought to alter the activity of additional channels and trigger further downstream signaling pathways (Nguyen et al., 2018; Farmer et al., 2020; Vega-Muñoz et al., 2020; Duong et al., 2021). In addition, the plant hormone jasmonic acid (JA) is produced at very high levels following the disruption of membranes and the oxidation of different fatty acids (Tripathi et al., 2017; Farmer and Goossens, 2019). JA, as well as other plant hormones, such as ethylene and salicylic acid, are also known to play a key role in the activation of wound-induced signaling.

In addition to triggering transcriptomic, metabolic, physiological and/or and proteomic responses at the tissues directly subjected to wounding (*i*.*e*., the wounded cells themselves, or cells neighboring the wounded cells; will be referred to here as ‘local tissue’), wounding is also accompanied by the triggering of systemic signal transduction pathways that activate wound-associated response mechanisms in tissues that have not been directly subjected to wounding (will be referred to here as ‘systemic tissue’). This process is termed systemic wound response (SWR) and is thought to enhance the overall resistance of plants to herbivory attack or other wound-related stresses (Walker-Simmons et al., 1984). Systemic signals thought to be involved in SWR include rapid changes in membrane potential (electric signals), calcium waves, ROS waves, and hormones such as JA (that can also function as volatiles; *e*.*g*., methyl JA), as well as other hormones and metabolites (*e*.*g*., Tripathi et al., 2017; Nguyen et al., 2018; Toyota et al., 2018; Fichman et al., 2019; Farmer et al., 2020; Matthus et al., 2020; Shao et al., 2020).

The release of glutamate from wounded cells, as well as local changes in pH, were recently proposed to play a key role in triggering SWRs by activating GLRs and initiating the systemic calcium wave (Toyota et al., 2018; Shao et al., 2020). In addition to the calcium wave, GLRs were previously shown to be required for the activation of systemic electric waves during wounding (Mousavi et al., 2013; Nguyen et al., 2018). A recent study showed that GLRs are also required for triggering the ROS wave during wounding (Fichman and Mittler, 2021). In contrast to glutamate, the role of eATP in triggering systemic responses to wounding is not established. Extracellular ATP was shown to trigger local ROS production during wounding following the sensing of eATP by the eATP receptor P2K that phosphorylates and activates the respiratory burst oxidase homolog D (RBOHD) protein (Chen et al., 2017). A new study has also shown that eATP is activating calcium signaling in cells via cyclic nucleotide gated ion channel 6 (CGNC6; Duong et al., 2021), and this process could also activate the calcium and ROS waves (Matthus et al., 2020).

The ROS wave is an auto-propagating cell-to-cell systemic signaling pathway triggered in plants in response to wounding and many different biotic and abiotic stresses (*e*.*g*., Fichman et al., 2019; Fichman and Mittler, 2020; Zandalinas et al., 2020a; Zandalinas et al., 2020b; Zandalinas and Mittler, 2021). It is dependent on the function of RBOHD and can travel in a cell-to-cell fashion over long distances from its site of initiation to the entire plant within minutes (Miller et al., 2009; Fichman et al., 2019; Fichman and Mittler, 2020). A recent study showed that the activation of the ROS wave in plants during wounding does not occur in mutants deficient in GLR (*i*.*e*., the *glr3*.*3glr3*.*6* double mutant), RBOHD (*rbohD*), or plasmodesmata (PD) functions (*i*.*e*., mutations in PD-localized protein 5; *pdlp5*) (Fichman and Mittler, 2021). However, the role of eATP in triggering the ROS wave and other systemic responses during wounding is not clear.

Because RBOHD plays a critical role in eATP-induced ROS formation during wounding (Chen et al., 2017), as well as in triggering and propagating the ROS wave (Miller et al., 2009; Fichman and Mittler, 2020), we used our newly developed whole-plant imaging system (Fichman et al., 2019; Zandalinas et al., 2020a; Zandalinas et al., 2020b; Fichman and Mittler, 2021; Zandalinas and Mittler, 2021) to elucidate the role of eATP in the wound-induced ROS wave response. Here we reveal that eATP that accumulates in wounded leaves and is sensed by P2K is required for the activation of the ROS wave during wounding. eATP application to unwounded leaves triggered the ROS wave, and the activation of the ROS wave was suppressed in mutants deficient in P2K (*i*.*e*., *p2k1-3, p2k2*, and *p2k1-3p2k2*). In addition, the expression of several SWR transcripts was suppressed in mutants deficient in P2K during wounding. Our findings reveal that in addition to sensing Glu via GLRs, eATP sensed by P2Ks is playing a key role in triggering the SWR of plants.

## RESULTS AND DISCUSSION

### Whole plant live imaging of eATP accumulation in plants following wounding

Extracellular ATP was previously imaged following wounding of plant tissues using a combination of luciferase and luciferin that produces light in the presence of ATP (*e*.*g*., Kim et al., 2006; Roux, 2014). These measurements were however restricted to cells and tissues and conducted using different microscopy platforms. To image eATP accumulation following wounding in whole *Arabidopsis thaliana* plants grown in soil, we used the same imaging platform we previously used for live imaging of ROS in whole plants in response to different stresses/treatments (*i*.*e*., the IVIS Lumina S5 platform; Fichman et al., 2019). We applied a mixture of luciferin and luciferase to plants and imaged light emission from plants following wounding. As shown in Figure 1, wounding of a single leaf of Arabidopsis resulted in the rapid accumulation of eATP (detected as early as 5 min following wounding; the earliest time point measured). The levels of eATP appear however to subside at 20 min following wounding. Systemic accumulation of eATP in non-wounded leaves of treated plants or in unwounded plants was not detected. The findings presented in Figure 1 demonstrate that wounding is followed by localized eATP accumulation, but that unlike ROS, the levels of eATP in systemic tissues does not change following wounding (at least within the time frame and detection limits of our method).

**Figure 1.**
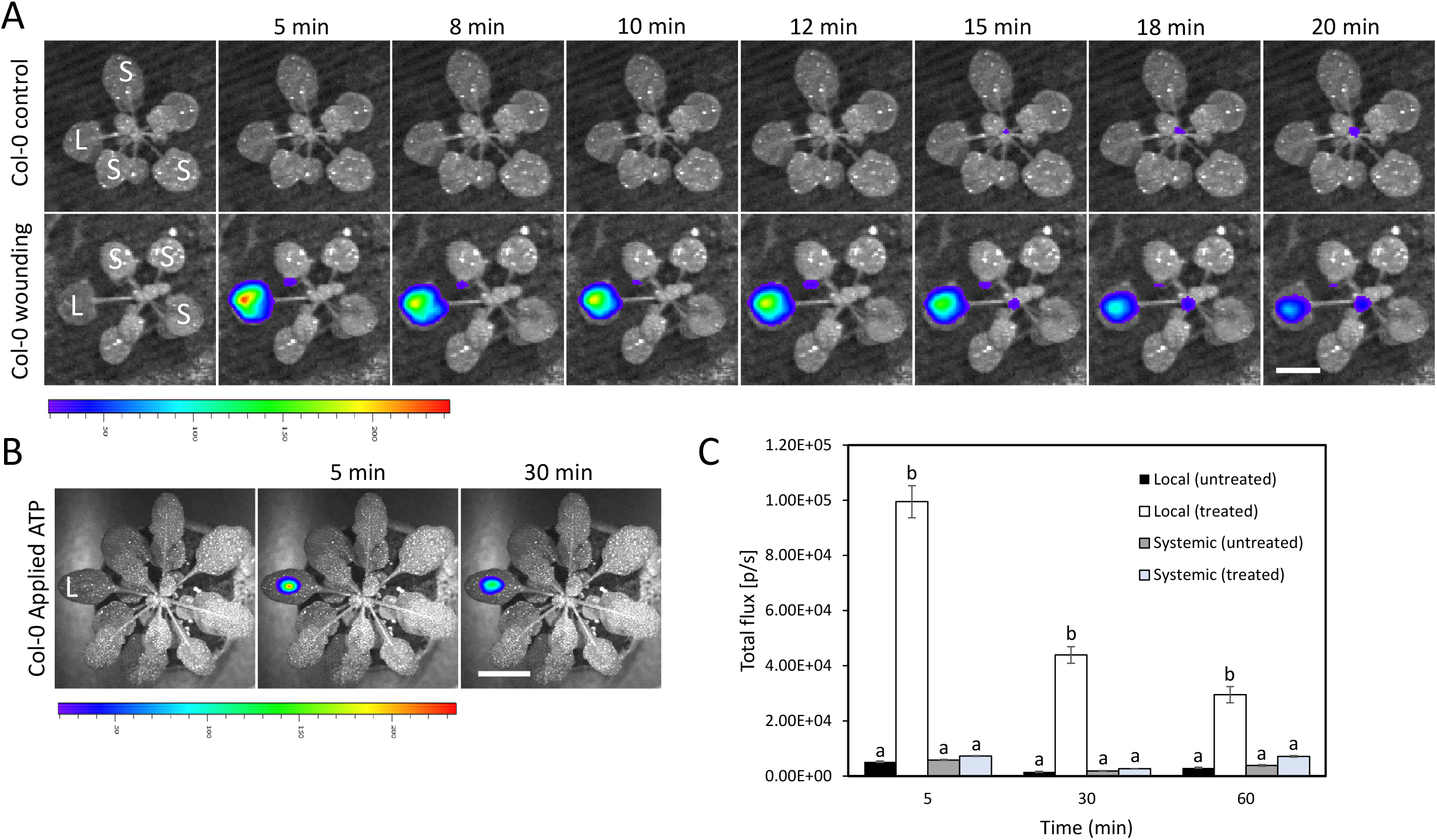
Whole plant live imaging of eATP accumulation following wounding. A, Representative time-lapse images of whole plant eATP levels (indicated by luciferase activity) in wild type Arabidopsis plants subjected to wounding on a single leaf (local; L). B, Representative time-lapse images of luciferase activity following application of a single drop of ATP (10 mM) to a single leaf (L; used as appositive control). C, Statistical analysis of eATP levels in local (L) and systemic (S) leaves of wounded (on leaf L) and unwounded plants at 5-, 30- and 60-min following wounding. All experiments were repeated at least three times with at least 3 treated and 3 untreated plants per repeat. Letters indicate statistical significance (P<0.05) by one-way ANOVA (n = 6). Bar = 1 cm. Abbreviations: eATP, extracellular ATP; L, local; S, systemic.

### Application of eATP to unwounded plants triggers the ROS wave

Because wounding was previously shown to trigger the ROS wave (Fichman et al., 2019), as well as to release ATP from cells (that becomes eATP; Figure 1; Kim et al., 2006; Roux, 2014), and the sensing of eATP by P2K was shown to trigger ROS production by RBOHD (Chen et al., 2017), we tested whether application of a stable form of eATP (*βγ*meATP; Chen et al., 2017; Pham et al., 2020; Duong et al., 2021) can trigger the ROS wave. As shown in Figure 2A, application of eATP to unwounded plants triggered the rapid accumulation of ROS in local and systemic leaves. This finding suggested that similar to different biotic and abiotic stresses such as heat, high light or pathogen infection, applied to a local leaf (*e*.*g*., (Fichman et al., 2019; Zandalinas et al., 2020a; Zandalinas et al., 2020b), eATP application is triggering the ROS wave. The activation and propagation of the ROS wave by different abiotic and biotic stimuli is dependent on the function of RBOHD that is thought to drive the accumulation of ROS at the apoplast (Miller et al., 2009; Fichman et al., 2019). To test whether the local and systemic accumulation of ROS by eATP is also dependent on RBOHD, we applied eATP to the *rbohD* mutant. As shown in Figure 2B, eATP application to the *rbohD* mutant did not result in local and systemic ROS accumulation. The findings presented in Figure 2 suggest that eATP application to plants can trigger the ROS wave.

**Figure 2.**
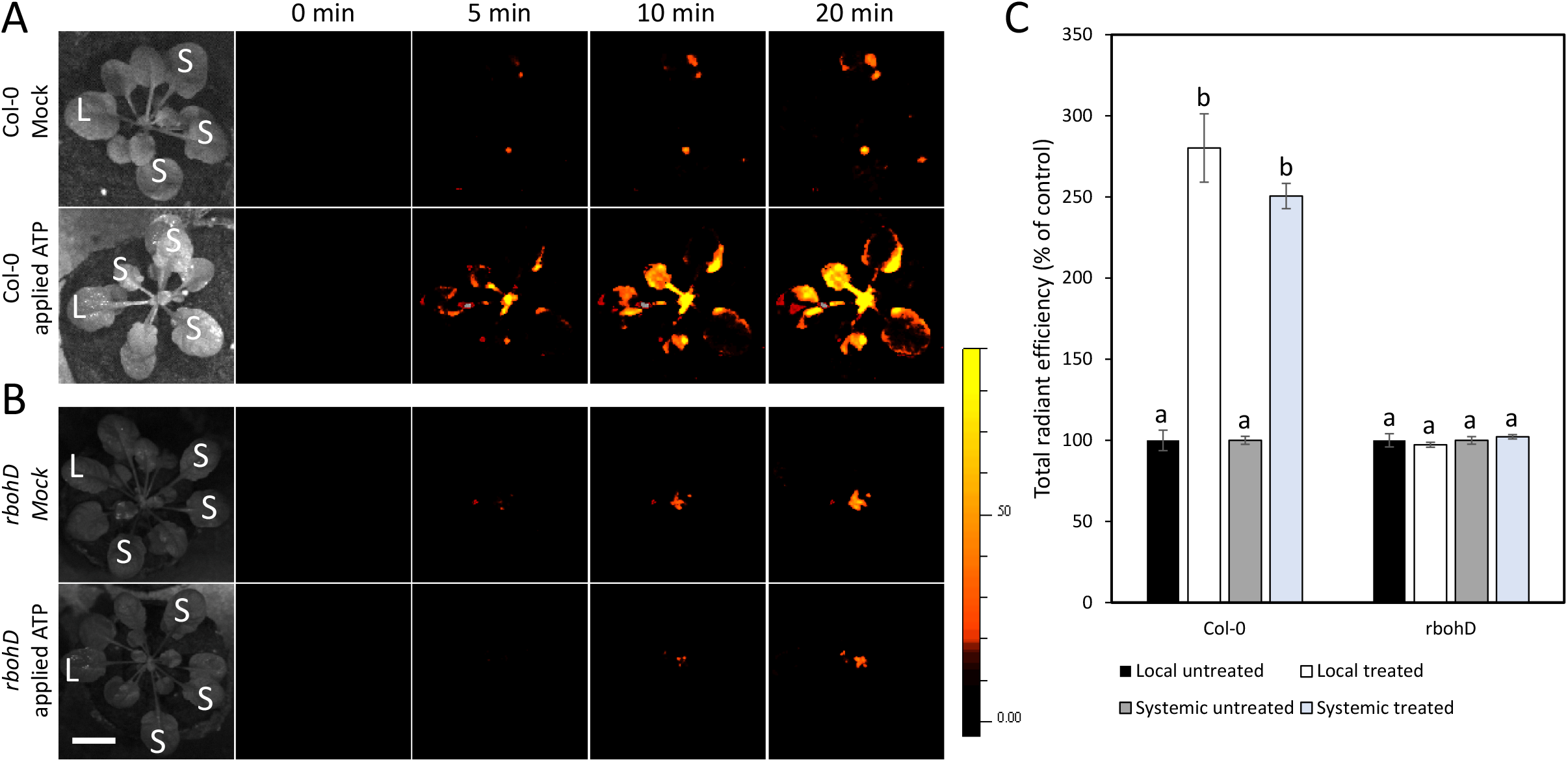
Application of eATP to unwounded plants triggers the ROS wave. A, Representative time-lapse images of whole-plant ROS levels (indicated by DCF oxidation) in wild type Arabidopsis plants following application of a single drop of *βγ*meATP (300 μM) to a single leaf (local; L). A single drop of buffer without ATP was applied to leaf L of mock untreated plants. B, Same as in A, but for *rbohD* mutants. C, Statistical analysis of local (L) and systemic (S) ROS levels in leaves of treated or mock untreated (on leaf L) wild type and *rbohD* plants at 20 min following buffer/eATP application. All experiments were repeated at least three times with at least 3 treated and 3 untreated plants per repeat. Letters indicate statistical significance (P<0.05) by one-way ANOVA (n = 6). Bar = 1 cm. Abbreviations: eATP, extracellular ATP; L, local; *rbohD*, respiratory burst oxidase homolog D; ROS, reactive oxygen species; S, systemic.

### The activation of the ROS wave following application of eATP is dependent on the eATP receptor P2K

Extracellular ATP is sensed in plant cells via the eATP receptor P2K leading to the activation of different metabolic and physiological responses (Choi et al., 2014; Chen et al., 2017; Pham et al., 2020). To test whether eATP application is triggering the ROS wave through the P2K pathway, we applied eATP to a single leaf of wild type (WT) and P2K mutants (*i*.*e*., *p2k1-3, p2k2*, and *p2k1-3p2k2*; Choi et al., 2014; Chen et al., 2017; Pham et al., 2020) and measured local and systemic ROS accumulation. As shown in Figure 3, compared to WT, the local and systemic accumulation of ROS in response to eATP was completely blocked in the *p2k1-3, p2k2*, and *p2k1-3p2k2* mutants. This finding suggests that eATP can trigger the ROS wave via the P2K receptors.

**Figure 3.**
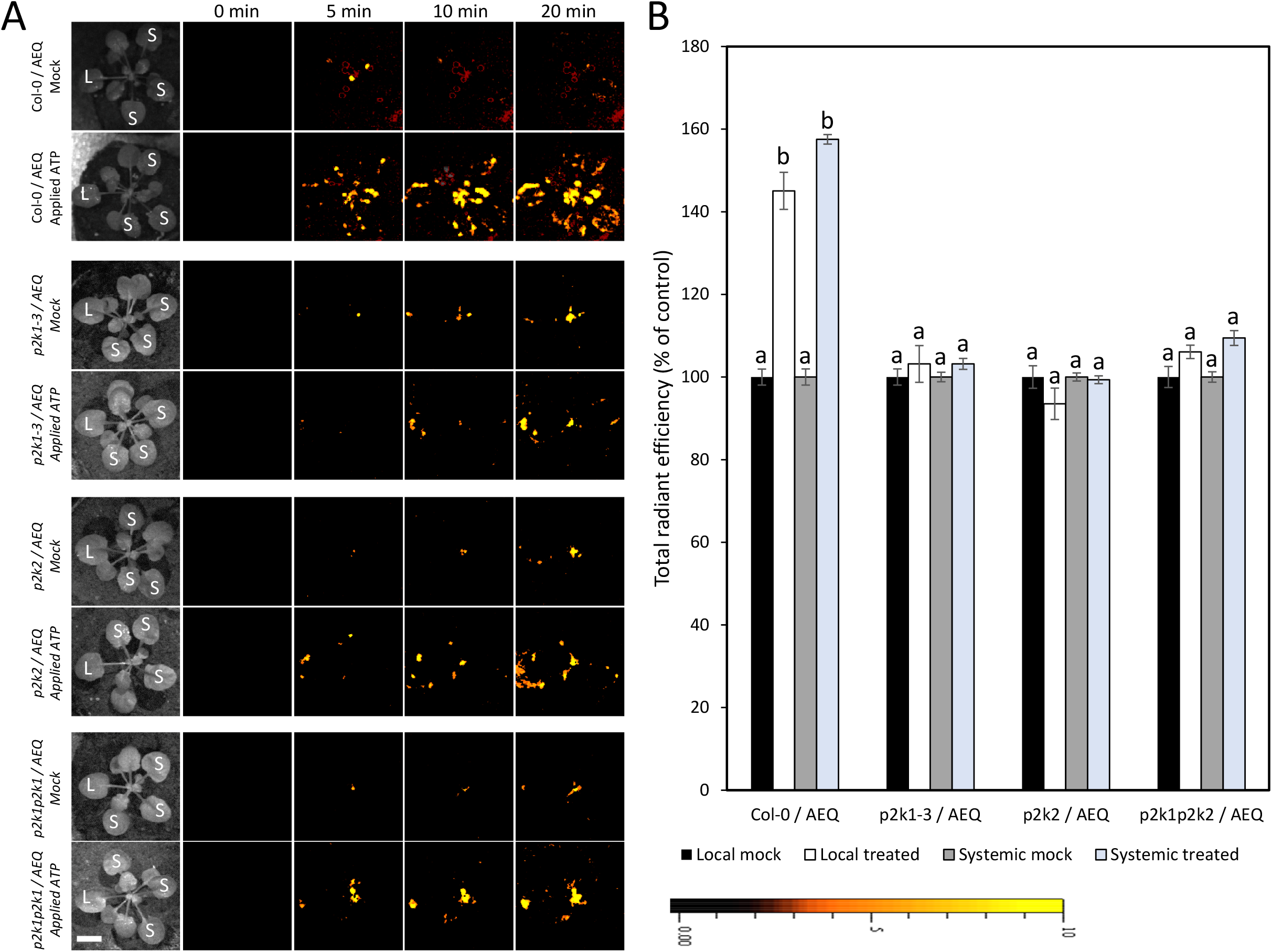
The activation of the ROS wave following application of eATP is dependent on the eATP receptor P2K. A, Representative time-lapse images of whole plant ROS levels (indicated by DCF oxidation) in wild type and *p2k1-3, p2k2*, and *p2k1-3p2k2* plants following application of a single drop of *βγ*meATP (300 μM) to a single leaf (local; L). A single drop of buffer without ATP was applied to leaf L of mock untreated plants. B, Statistical analysis of local (L) and systemic (S) ROS levels in leaves of treated or mock untreated (on leaf L) wild type and *p2k1-3, p2k2*, and *p2k1-3p2k2* mutant plants at 20 min following buffer/eATP application. All experiments were repeated at least three times with at least 3 treated and 3 untreated plants per repeat. Letters indicate statistical significance (P<0.05) by one-way ANOVA (n = 6). Bar = 1 cm. Abbreviations: eATP, extracellular ATP; L, local; P2K, purinoreceptor 2 kinase; ROS, reactive oxygen species; S, systemic.

### The activation of the ROS wave following wounding is suppressed in mutants deficient in the eATP receptor P2K

The sensing of eATP by the eATP receptor protein P2K was previously shown to trigger ROS production in cells via phosphorylation of RBOHD (Chen et al., 2017). To test whether P2K is also involved in the activation of the ROS wave during wounding (Miller et al., 2009; Fichman et al., 2019; Fichman and Mittler, 2021), we measured local and systemic ROS accumulation in the *p2k1-3, p2k2*, and *p2k1-3p2k2* mutants. As shown in Figure 4, compared to WT, the accumulation of local and systemic ROS in the P2K mutants was suppressed following wounding of a local leaf. These finding suggest that the sensing of eATP during wounding plays an important role in triggering the ROS wave. The finding that in contrast to eATP application (Figure 3), the ROS wave was not completely suppressed in the P2K mutants following wounding (Figure 4) suggests however that additional wound-related signals, for example Glu (Mousavi et al., 2013; Toyota et al., 2018; Shao et al., 2020), could also be involved in the triggering of the ROS wave following mechanical wounding. Indeed, the ROS wave was previously shown to be suppressed in the *glr3*.*3glr3*.*6* double mutant in response to wounding (Fichman and Mittler, 2021).

**Figure 4.**
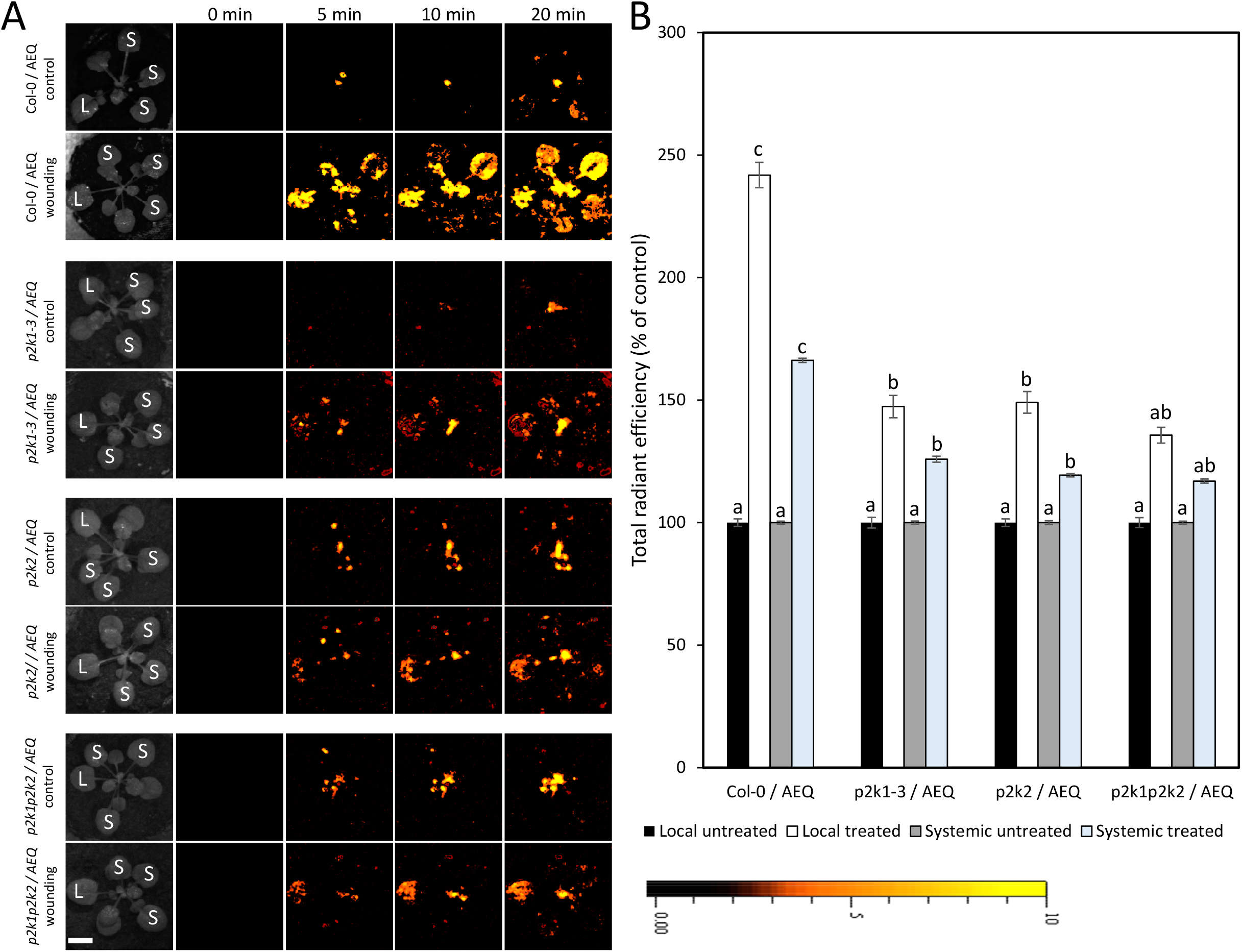
The activation of the ROS wave following wounding is suppressed in mutants deficient in the eATP receptor P2K. A, Representative time-lapse images of whole plant ROS levels (indicated by DCF oxidation) in wild type and *p2k1-3, p2k2*, and *p2k1-3p2k2* plants following wounding of a single leaf (local; L). B, Statistical analysis of local (L) and systemic (S) ROS levels in leaves of wounded or unwounded (on leaf L) wild type and *p2k1-3, p2k2*, and *p2k1-3p2k2* mutant plants at 20 min following wounding. All experiments were repeated at least three times with at least 3 treated and 3 untreated plants per repeat. Letters indicate statistical significance (P<0.05) by one-way ANOVA (n = 6). Bar = 1 cm. Abbreviations: eATP, extracellular ATP; L, local; P2K, purinoreceptor 2 kinase; ROS, reactive oxygen species; S, systemic.

### The expression of wound- and ROS-response transcripts is suppressed in systemic leaves of mutants deficient in the eATP receptor P2K following local wounding

The ROS wave was previously shown to be required for systemic transcriptomic responses to local heat, high light, or mechanical stress treatments (Suzuki et al., 2013; Fichman et al., 2020; Zandalinas et al., 2020a; Zandalinas et al., 2020b; Fichman and Mittler, 2021). However, the role of eATP, P2K and the ROS wave in triggering systemic transcriptomic responses to wounding is not clear. Because the ROS wave is suppressed following wounding in the P2K mutants (Figure 4), we tested whether this suppression is also affecting the expression of several wound- and ROS-response systemic transcripts (*i*.*e*., *ZAT10, ZAT12* and *WRKY40*; Suzuki et al., 2013; Zandalinas et al., 2019; Zandalinas et al., 2020a; Zandalinas et al., 2020b; Fichman and Mittler, 2021). As shown in Figure 5, compared to WT, the local and systemic expression of *ZAT10* and *WRKY40*, and the systemic expression of *ZAT12*, were suppressed in the *p2k1-3, p2k2*, and *p2k1-3p2k2* mutants. This finding suggests that the sensing of eATP plays an important role not only in the triggering of the ROS wave following wounding, but also in the activation of some transcriptomic responses in systemic leaves.

**Figure 5.**
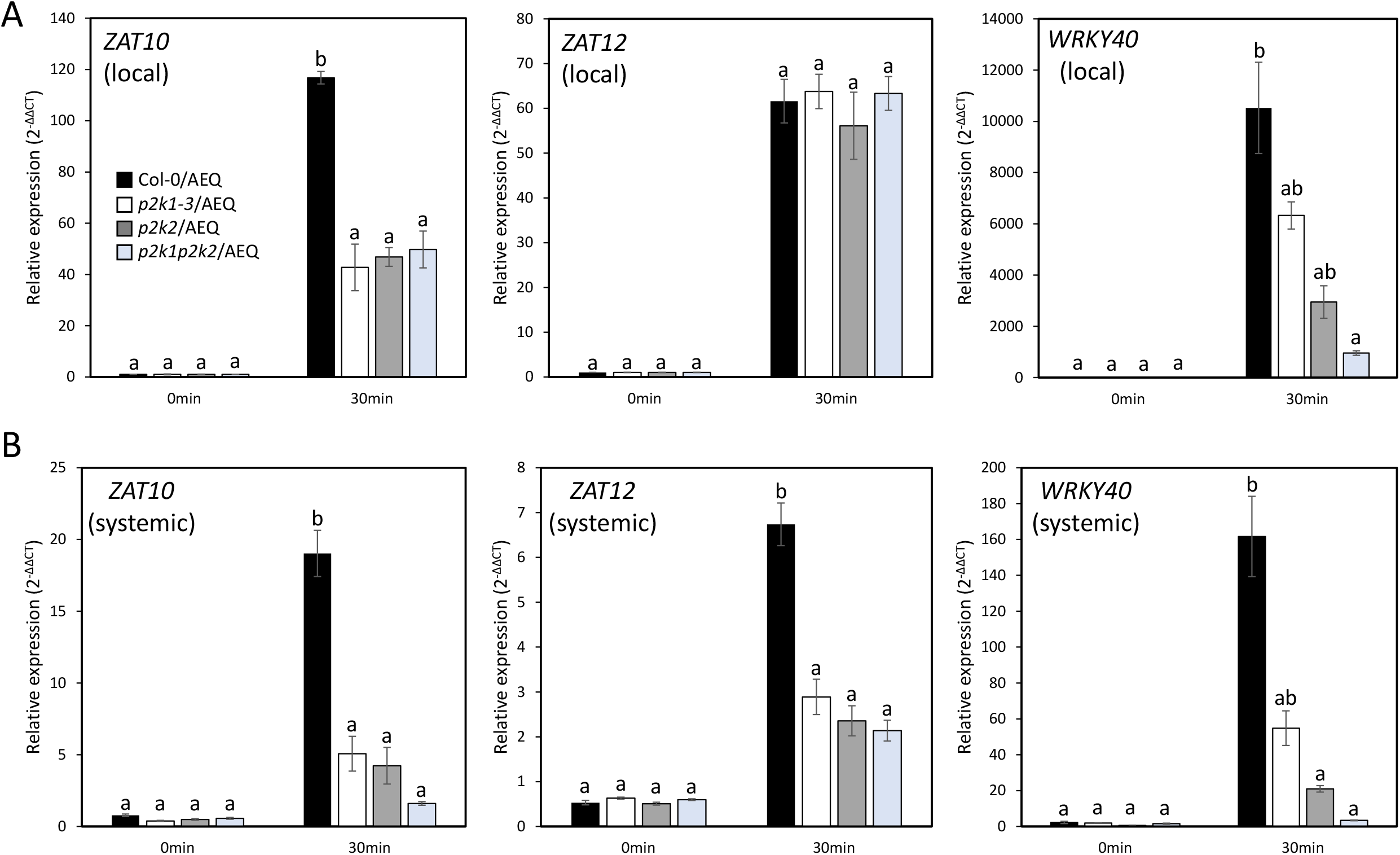
The expression of wound- and ROS-response transcripts is suppressed in systemic leaves of mutants deficient in the eATP receptor P2K following local wounding. A, Real-time quantitative polymerase chain reaction (RT-qPCR) analysis for *ZAT10, ZAT12*, and *WRKY40* steady-state transcript levels in local leaves of wild type and *p2k1-3, p2k2*, and *p2k1-3p2k2* plants following wounding of a single leaf. B, Same as in A, but for systemic leaves. Transcript expression is represented as the relative quantity (2^−ΔΔCT^) compared to an internal control (elongation factor 1α) in unwounded local tissue of wild-type (time 0). All experiments were repeated at least three times with at least 3 treated and 3 untreated plants per repeat. Letters indicate statistical significance (P<0.05) by one-way ANOVA (n = 6). Abbreviations: eATP, extracellular ATP; L, local; P2K, purinoreceptor 2 kinase; S, systemic.

### Application of Glu, eATP, or eATP+Glu to unwounded plants triggers the ROS wave

To test whether eATP and Glu could have a synergistic effect on the activation of the ROS wave, we applied eATP, Glu, or eATP+Glu to a local leaf and measured the local and systemic accumulation of ROS. As shown in Figure 6A, application of Glu, eATP, or eATP+Glu resulted in a similar systemic ROS accumulation response. In contrast, compared to eATP, application of Glu to a local leaf generated a lower local ROS response. These finding suggest that Glu or eATP can trigger the ROS wave during wounding, but that they likely trigger it via the same pathway.

**Figure 6.**
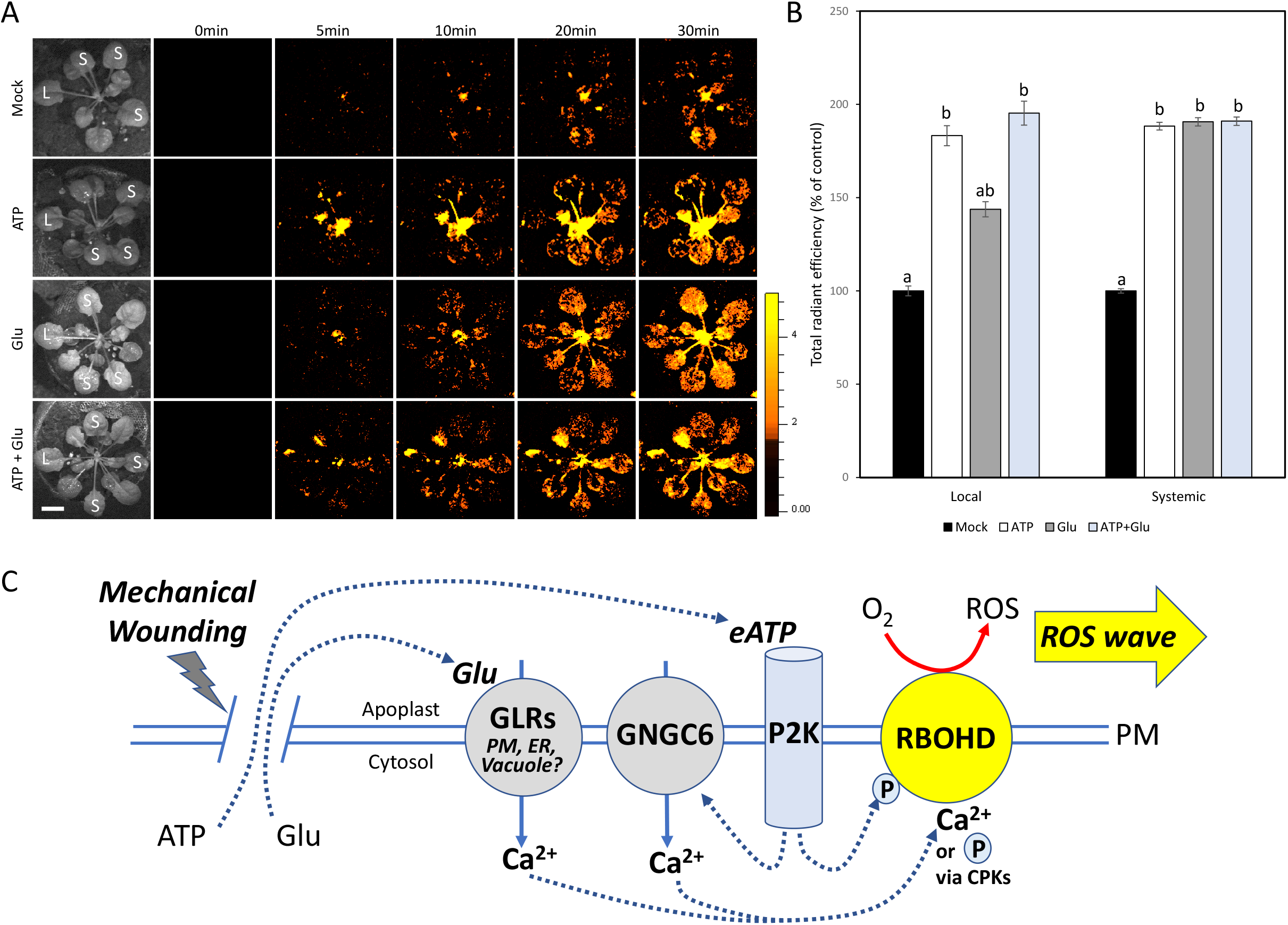
Application of Glu, eATP, or eATP+Glu to unwounded plants triggers the ROS wave and a model. A, Representative time-lapse images of whole-plant ROS levels (indicated by DCF oxidation) in wild type Arabidopsis plants following application of a single drop of *βγ*meATP (300 μM), Glu (50 mM), or *βγ*meATP (300 μM) + Glu (50 mM) to a single leaf (local; L). A single drop of buffer without ATP or Glu was applied to leaf L of mock untreated plants. B, Statistical analysis of local (L) and systemic (S) ROS levels in leaves of treated or mock untreated (on leaf L) plants at 20 min following buffer/eATP/Glu/ATP+Glu application. All experiments were repeated at least three times with at least 3 treated and 3 untreated plants per repeat. Letters indicate statistical significance (P<0.05) by one-way ANOVA (n = 6). Bar = 1 cm. C, A hypothetical model depicting the proposed role of eATP in triggering the ROS wave during wounding. The sensing of eATP by P2Ks, as well as the sensing of Glu by GLRs is shown to trigger ROS production by RBOHD and initiate the ROS wave. Abbreviations: eATP, extracellular ATP; CNGC, cyclic nucleotide-gated channel; CPK, calcium-dependent kinase; ER, endoplasmic reticulum; GLR, glutamate-like receptor; Glu, glutamate; L, local; P, phosphate; P2K, purinoreceptor 2 kinase; PM, plasma membrane; rbohD, respiratory burst oxidase homolog D; ROS, reactive oxygen species; S, systemic.

## CONCLUSIONS

The local wound response is a complex response that involves the activation of many different signal transduction pathways. These include pathways responding to membrane depolarization, eATP, Glu, pH, ROS, and different plant hormones, including JA (*e*.*g*., Mousavi et al., 2013; Tripathi et al., 2017; Nguyen et al., 2018; Toyota et al., 2018; Farmer et al., 2020; Ikeuchi et al., 2020; Shao et al., 2020; Vega-Muñoz et al., 2020; Moore et al., 2021). While Glu, pH, and membrane depolarization were recently shown to trigger the SWR via GLR-triggered electric, calcium and ROS waves (Mousavi et al., 2013; Nguyen et al., 2018; Toyota et al., 2018; Fichman and Mittler, 2021; Figure 6A), the role of eATP in triggering SWR is not established. Our study reveals that during mechanical wounding eATP plays an important role in triggering the ROS wave and SWRs, and that eATP triggers these responses via the P2K-RBOHD pathway (Chen et al., 2017; Figure 6B). However, because the ROS wave and the SWR are not completely blocked in the P2K mutants (Figures 4 and 5), it is likely that other pathways involved in SWR (*e*.*g*., the Glu-GLR pathway; Mousavi et al., 2013; Nguyen et al., 2018; Toyota et al., 2018; Shao et al., 2020; Figure 6A) could be playing a role in this response (Figure 6B). Further studies are therefore needed to determine how eATP is integrated with other wound-induced signal transduction pathways. When it comes to local and systemic ROS production during wounding, our findings that RBOHD is essential for ROS accumulation in response to eATP (Figure 2), as well as in response to wounding (Fichman and Mittler, 2021), suggests that RBOHD is a central integrator of wound-induced ROS associated signals (Figure 6B). RBOHD could integrate eATP-driven P2K phosphorylation, with GLR- and CNGC-driven calcium, and/or GLR- and CNGC-driven calcium dependent kinase (CPKs; *e*.*g*., Dubiella et al., 2013) signals to trigger the ROS wave (Figure 6B).

## MATARIALS AND METHODS

### Plant Materials, Growth Conditions, and Stress Treatments

*Arabidopsis thaliana* (Col-0) wild-type, *rbohD* (AT5G47910; Torres et al., 2002), and aequorin-expressing wild-type and *p2k1-3* (SALK_042209; Choi et al., 2014; Chen et al., 2017), *p2k2* (GK-777H06; Pham et al., 2020), and *p2k1p2k2* (SALK_042209/GK-777H06; Pham et al., 2020) plants were grown on peat pellets (Jiffy 7; Jiffy International) for 4 weeks under controlled short-day conditions of 10-h-light/14-h-dark, 50 μmol m^-2^s^-1^, and 21°C ambient temperature. Wounding was applied to a single leaf by puncturing with 18 dressmaker pins simultaneously as described in Fichman et al., 2019. Extracellular ATP was applied to a single leaf via submersion for 5 seconds in dipping solution of 300 μM *βγ*meATP (Millipore-Sigma), 50 μM H_2_DCFDA (Millipore-Sigma), and 2.67 mM MgSO_4_ (Fisher) in 0.1 mM EDTA, 0.05 M phosphate buffer, pH 7.4, 0.01% (v/v) Silwet L-77, as described in Kim et al., 2006 and Fichman et al., 2019.

### Extracellular ATP Imaging

To visualize extracellular ATP following wounding, plants were sprayed until leaf saturation and runoff with 1 mM luciferin (sodium salt; GOLD Biotechnology; Miller et al., 2009), 10 μg / mL firefly luciferase (Promega), and 5mM MgCl_2_ in 0.5M Tris-HCl, pH 7.5. A local leaf was then immediately wounded as described above. After an initial 5 min incubation period in darkness (to eliminate light-induced bioluminescent background), images of the resulting bioluminescence were acquired for 1 hr using the IVIS Lumina S5 system (PerkinElmer) and analyzed using Living Image 4.7.2 software as described in Fichman et al., 2019 and Zandalinas et al., 2020b. Quantification of bioluminescence in regions of interest was calculated using counts (photons/second) as described by Zandalinas et al., 2020b. Positive bioluminescent controls were performed by the application of a drop of ATP (10mM; Millipore-Sigma) to a single leaf with no wounding. All experiments were repeated at least three times with at least three biological repeats per treatment.

### ROS Imaging

Whole-plant ROS accumulation was imaged and analyzed as described in Fichman et al., 2019. Plants were fumigated for 30 minutes with 50 μM H_2_DCFDA (Millipore-Sigma) in 0.05M phosphate buffer, pH 7.4, 0.01% (v/v) Silwet L-77 using a nebulizer (Punasi Direct) within a glass enclosure to allow for uptake of the vaporized solution into the plants. Following fumigation, a single leaf was wounded or treated with eATP as described above and images of the resulting ROS wave were captured over 30 minutes using the IVIS Lumina S5 system (PerkinElmer). Images were analyzed using Living Image 4.7.2 and data analysis was performed by measuring radiant efficiency in regions of interest as described in Fichman et al., 2019. To ensure penetration of the H_2_DCFDA dye into plants, controls were performed by fumigation with 0.3% (v/v) H_2_O_2_ for 10 min after 50 μM H_2_DCFDA fumigation for 30 min and image acquisition in the IVIS Lumina S5 system (Fichman et al., 2019; Fichman and Mittler, 2020; Zandalinas et al., 2020b). All experiments were repeated at least three times with at least three biological repeats per treatment.

### Transcript Expression Analysis

Transcriptional responses of local and systemic leaves following wounding were analyzed at 0- and 30-min following wounding. Young systemic leaves located approximately 137.5° away from the sampled local leaf within the rosette were chosen for sampling. Plants wounded at a single leaf were allowed to incubate for 30 min following wounding prior to sampling of the local and systemic leaf. Four biological repeats for each treatment were used each with 12 technical repeats. RNA extraction was performed using Plant RNeasy kit (Qiagen) as described by the manufacturer. Complementary DNA was synthesized from the quantified extracted RNA template (Primescript RT Reagent Kit, Takara Bio). Real-time quantitative polymerase chain reaction (RT-qPCR) analysis of transcript expression was performed for genes *ZAT10* (AT5G59820), *ZAT12* (AT1G27730), and *WRKY40* (AT1G80840) using iQ SYBR Green supermix (Bio-Rad) in the CFX Connect Real-Time PCR Detection System (Bio-Rad) as described in Fichman and Mittler, 2021. *ELONGATION FACTOR 1A* served as the internal control to normalize relative gene expression across samples. Primer sequences used for each transcript are shown in Supplementary Table S1. Quantification of normalized relative gene expression (2^-ΔΔCT^) is presented as percent of control (with the unwounded local sample acting as the control).

### Statistical Analysis

Statistical analysis was performed with one-way ANOVA followed by the Tukey post hoc test. Results are shown as the means for the data, with ± SE indicated for each value. Letters represent a statistically significant difference corresponding to at least P<0.05.

---

**Supplemental Table S1.**
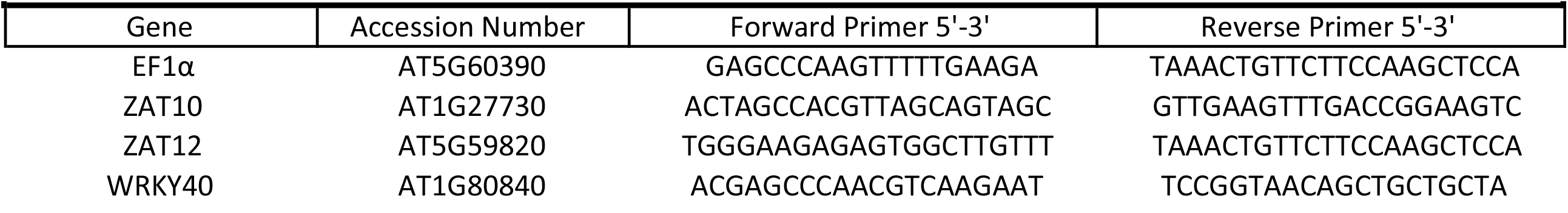
Primers for transcript expression analysis via RT-qPCR.

